# Optimizing complex phenotypes through model-guided multiplex genome engineering

**DOI:** 10.1101/086595

**Authors:** Gleb Kuznetsov, Daniel B. Goodman, Gabriel T. Filsinger, Matthieu Landon, Nadin Rohland, John Aach, Marc J. Lajoie, George M. Church

**Author notes:** These authors contributed equally to this work (Gleb Kuznetsov, Daniel B. Goodman, Gabriel T. Filsinger). Correspondence should be addressed to M.J.L. or G.M.C.

## Abstract

Optimization of complex phenotypes in engineered microbial strains has traditionally been accomplished by laboratory evolution. However, only a subset of the resulting mutations may affect the phenotype of interest and many others may have unintended effects. Multiplexed genome editing can complement evolutionary approaches by creating diverse combinations of targeted changes, but in both cases it remains challenging to identify which alleles influence the desired phenotype. We present a method for identifying a minimal set of genomic modifications that optimizes a complex phenotype by combining iterative cycles of multiplex genome engineering and predictive modeling. We applied our method to the 63-codon *E. coli* strain C321.ΔA, which has 676 mutations relative to its wild-type ancestor, and identified six single nucleotide mutations that together recover 59% of the fitness defect exhibited by the strain. The resulting optimized strain, C321.DA.opt, is an improved chassis for production of proteins containing non-standard amino acids. Our data reveal how multiple cycles of multiplex automated genome engineering (MAGE) and inexpensive sequencing can generate rich genotypic and phenotypic diversity that can be combined with linear regression techniques to quantify individual allelic effects. While laboratory evolution relies on enrichment as a proxy for allelic effect, our model-guided approach is less susceptible than enrichment to bias from population dynamics and recombination efficiency. We also show that the method can identify beneficial *de novo* mutations that arise adventitiously. Beyond improving the fitness of C321, ΔA, our work provides a proof-of-principle for high-throughput quantification of individual allelic effects which can be used with any method for generating targeted genotypic diversity.

## Background

Genome editing and DNA synthesis technologies are enabling the construction of engineered organisms with synthetic metabolic pathways ^1^, reduced and refactored genomes ^2–5^, and expanded genetic codes ^6,7^. However, genome-scale engineering can come at the cost of reduced fitness or suboptimal traits ^2,6,7^ caused by design flaws that fail to preserve biologically-important features ^7,8^, synthesis errors, or collateral mutations acquired during strain construction ^6^. It remains challenging to identify alleles that contribute to these complex phenotypes and it becomes prohibitive to test them individually. Laboratory evolution has traditionally been used to improve desired phenotypes and navigate genetic landscapes ^9^, but this process relies on mutations that accumulate across the genome and may disrupt synthetic designs or traits not maintained under selection. In contrast, targeted genome engineering can alter the genome at chosen loci and can be used to target many locations simultaneously ^12^. Multiplexed editing creates a large pool of combinatorial genomic changes than can be screened or selected to find high-performing genomic designs. However, as the number of targeted loci considered increases, it becomes difficult to interpret the significance of individual changes. There remains a need for a method to rapidly identify subsets of beneficial alleles from a large list of candidates and optimize large-scale genome engineering efforts.

Leveraging recent improvements in the cost and speed of microbial whole genome sequencing, we present a method for identifying precise genomic changes that optimize complex phenotypes, combining multiplex genome engineering, genotyping, and predictive modeling (**Fig. 1**). Multiple rounds of genome editing are used to generate a population enriched with combinatorial diversity at the targeted loci. Throughout the editing process, clones from the population are subject to whole-genome sequencing and are screened for phenotype. The genotype and phenotype data is used to update a model which predicts the effects of individual alleles. These steps are repeated on a reduced set of candidate alleles informed by the model, or on a new set of targets. Finally, the highest impact alleles are rationally introduced into the original organism, minimizing alterations to the organism’s original genotype while optimizing the desired phenotype.

**Figure 1.**
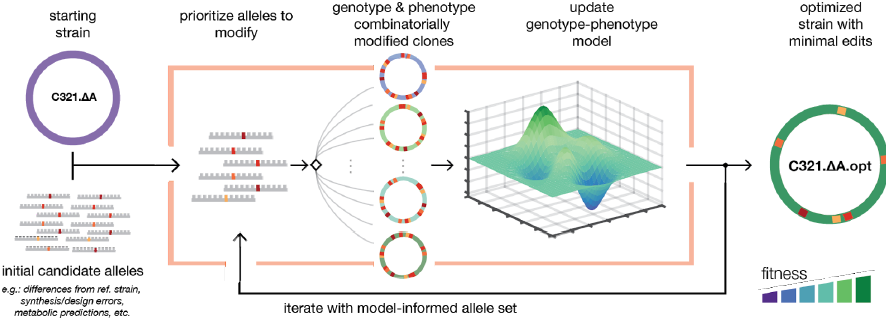
Workflow for improving phenotypes through model-guided multiplex genome editing. First, an initial set of target alleles (hundreds to thousands) is chosen for testing based on starting hypotheses. These targets may be designed based on differences from a reference strain, synthesis or design errors, or biophysical modeling. Multiplex genome editing creates a set of modified clones enriched with combinations of the targeted changes. Clones are screened for genotype and phenotype, and predictive modeling is used to quantify allele effects. The workflow is repeated to validate and test new alleles. Beneficial alleles are combined to create an optimized genotype.

We applied this method to the genomically recoded organism (GRO) C321.ΔA, a strain of *E*. *coli* engineered for non-standard amino acid (nsAA) incorporation ^6^. This GRO was constructed by replacing all 321 UAG stop codons with synonymous UAA codons and deleting UAG-mediated release factor *prfA*. Over the course of the construction process, C321.ΔA acquired 355 off-target mutations and developed a 60% greater doubling time relative to its non-recoded parent strain, *E*. *coli* MG1655. An improved C321.ΔA strain would accelerate the pace of research involving GROs and further enable applications leveraging expanded genetic codes, including biocontainment ^13^, virus resistance ^6^, ^14^ and expanded protein properties ^15^. We expected that a subset of the off-target mutations caused a significant fraction of the fitness defect, providing a starting hypothesis for iterative improvement.

## Results

To select an initial set of candidate alleles, we first used the genome engineering and analysis software *Millstone* (Goodman et al., submitted) to analyze sequencing data from C321.ΔA and identify all mutations relative to the parental strain MG1655. Millstone uses SnpEff ^16^ to annotate affected genes and predicted severity for each mutation. We further annotated each coding mutation with its associated growth defect in LB medium upon knockout of the affected gene after 22 hours (LB_22), as reported in the Keio collection ^17^. Based on this analysis, we identified 127 mutations in proteins and non-coding RNA as the top candidates responsible for fitness impairment. Our candidate alleles included all frameshift and non-synonymous mutations, mutations in non-coding RNA, and synonymous changes in genes with LB_22 < 0.7. We partitioned the targets into three priority categories according to predicted effect (**Supplementary Table 1** and **Supplementary Table 2**).

MAGE introduces combinations of genome edits with approximately 10-20% of cells receiving at least one edit per cycle ^12^. To generate a diverse population of mutants enriched for reversions at multiple loci, we performed up to 50 cycles of MAGE in three lineages. The first lineage used a pool of 26 oligonucleotides targeting only the highest category of mutations, the second lineage targeted the top 49 sites, and the third lineage targeted all 127 (**Supplementary Fig. 1**).

We sampled a total of 87 clones from multiple time points and lineages during MAGE cycling. We also sequenced three separate clones of the starting strain. We then performed whole genome sequencing and measured doubling time for each clone. *Millstone* was used to process sequencing data and to report variants for all 90 samples in parallel. We observed fitness improvement across all three lineages with a diversity of genotypes and fitness phenotypes across the multiple time points (**Fig. 2** and **Fig. 3a,b**). Clones selected from the final time point recovered 40-58% (mean 49%) of the fitness defect compared to MG1655 and had between 5 and 15 (mean 10.2) successfully reverted mutations. Of the 127 targeted mutations, 99 were observed in at least one clone, with as many as 19 successful reversions in a clone from the 127-oligo lineage. Additionally, we observed 1,329 unique de novo mutations across all clones (although only 135 were called in more than one clone), accumulating at a rate of roughly one per MAGE cycle in each clone (**Fig. 2d,e**). This elevated mutation rate was caused by defective mismatch repair (*mutS*-), which both increases MAGE allele replacement frequency and provides a source of new mutations that could improve fitness.

**Figure 2.**
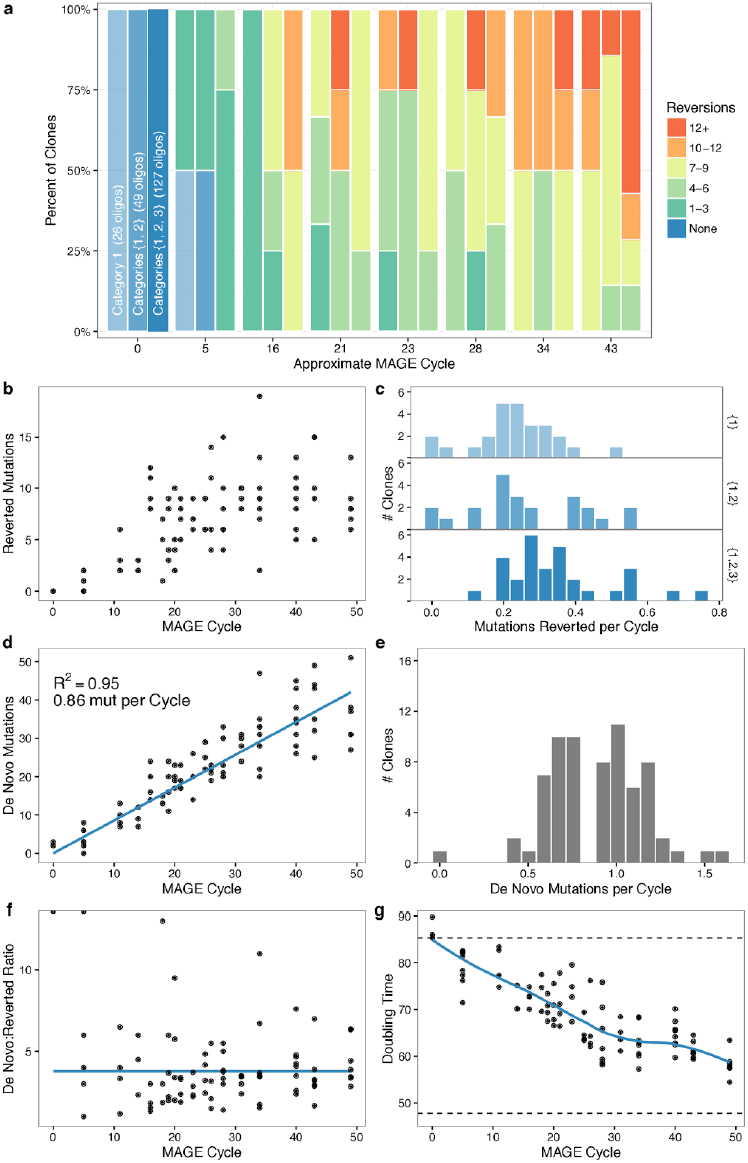
Mutation dynamics over many cycles of MAGE allele reversion. (**a**) Increase in combinatorial diversity and reversion count versus number of MAGE cycles. (**b**) Number of reversions per clone vs MAGE cycle. (**c**) The rate of reversions per MAGE cycle among the different allele categories, showing a higher rate per cycle for cells exposed to all 127 oligos. (**d**) The number of *de novo* mutations per clone over successive MAGE cycles. (**e**) Rate of de novo mutations per MAGE cycle. (**f**) The average ratio between number of de *novo* mutations and reverted alleles per MAGE cycle remains constant throughout the experiment. (**g**) Doubling time (min) improvement per clone from the C321.ΔA starting strain (top dotted line) towards the ECNR2 parent strain (bottom dotted line). Blue line is a LOESS fit.

**Figure 3.**
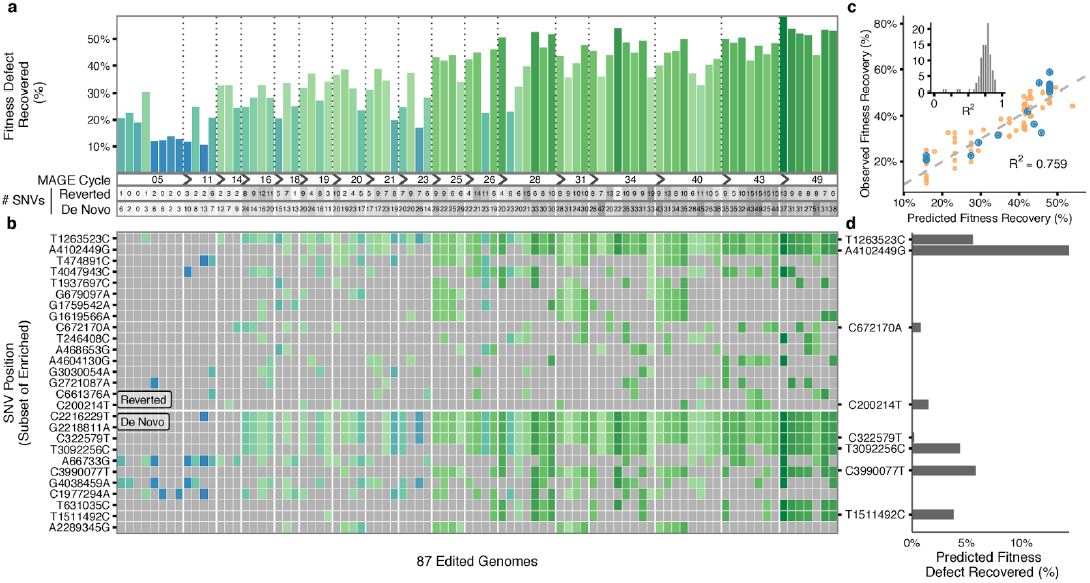
Genotypic and phenotypic diversity in 87 clones sampled across 50 MAGE cycles enabled model-guided prioritization of top single nucleotide variants (SNVs) for further validation. (**a**) Percent of C321.ΔA fitness defect recovered across MAGE cycles (shown with bar color and height). The number of SNVs reverted or introduced are shown below. (**b**) Presence of targeted reversions and de novo mutations in each clone colored according to fitness. A subset of the most enriched mutations are shown, ordered by enrichment (full dataset available in **Supplementary Table 9**). (**c**) Example model fit using top 8 alleles as features with 15 samples left out as a test set (blue points) and used to evaluate R^2^. Training points are plotted in orange. The inset shows distribution of R 2 values for 100 different simulations with 15 random samples left out to calculate R^2^ for each. Example fit was chosen to exemplify a median R^2^ value from this distribution. (**d**) Average model fit coefficients for top 8 alleles assigned non-zero values over repeated cross-validated linear regression (**Online Methods**) indicates their predicted contribution to fitness improvement.

The combinatorial diversity produced by multiplex genome engineering generates a dataset well-suited for analysis by linear regression. We made a simplifying assumption that doubling time is determined by the independent effects of individual alleles and employed a first-order multiplicative model that predicts doubling time based on allele occurrence (**Online Methods and Supplementary Note 1**). As model features, we considered the 99 reversions and 135 de novo mutations that occurred in at least two clones. Multiple linear regression was used to fit the model, with feature coefficients indicating the predicted effect of the respective allele. We expected a small number of alleles to contribute significantly to fitness improvement and thus used elastic net regularization 18 to impose this sparsity constraint on regression while accounting for high levels of co-occurrence among some alleles. To limit overfitting, we performed multiple rounds of k-fold cross-validation (k=5) and selected alleles that were assigned a non-zero coefficient on average. The analysis of the data obtained over 50 cycles of MAGE identified four targeted reversions and four de novo mutations that had the greatest putative effect on fitness (**Fig 3c,d** and **Supplementary Table 3).**

To validate the eight alleles prioritized in the 50-cycle MAGE experiment, we performed nine cycles of MAGE using a pool of eight oligos (**Supplementary Table 3**) applied to the starting C321.ΔA strain. We then screened each clone using MASC-PCR and measured doubling time (**Supplementary Fig. 2**). Modeling revealed strong effects for two reversions (T1263523C and A4102449G) and one de novo mutation (C3990077T), along with weaker effects for two additional reversions (C200214T and C672170A). These mutations are discussed in Supplementary Note 2. A clone with all five of these mutations was isolated and measured to have recovered 51% of the fitness defect in C321.ΔA. The three remaining de novo mutations did not show evidence of improving fitness despite being highlighted in the initial modeling, illustrating the importance of subsequent validation of model-selected alleles.

To identify mutations that further improved the fitness of C321, ΔA, we extended our search to off-target mutations occurring in regulatory regions using smaller pool sizes. We identified seven non-coding mutations predicted to disrupt gene regulation ^8^ (**Online Methods** and **Supplementary Table 4**). Applying nine rounds of MAGE followed by linear modeling revealed that reverting a single mutation in the −35 box of the folA promoter recovers a predicted 27% of the fitness defect (**Supplementary Fig. 3**). To test whether any of the designed UAG-to-UAA mutations caused a fitness defect in the C321 background, we followed the same procedure with 20 previously recoded UAA codons predicted to have a potential disruptive effect. (**Supplementary Table 5**). We tested reversion back to UAG in a *prfA*^+^ variant of C321 capable of terminating translation at UAG codons. We observed no evidence of a beneficial fitness effect from any individual UAA-to-UAG reversion.

Finally, we used MAGE to introduce the best six mutations (**Supplementary Table 6**) into the original C321.ΔA strain (**Online Methods**), creating an optimized strain C321.ΔA.opt that restores 59 +/-11% of the fitness defect in C321, ΔA (**Fig. 4a**). This rationally designed strain recovered the same amount of fitness as the fastest clones obtained through 50 rounds of MAGE and substantial passaging, which resulted in 6-13 reversions and 31-38 *de novo* mutations. (**Fig. 4a**). Whole genome sequencing of the final strain confirmed that no UAG codons were reintroduced. Nine additional *de novo* mutations arose, but these are predicted to have a neutral effect (**Supplementary Table 7**). We characterized UAG-dependent incorporation of the nsAAs p-acetyl-L-phenylalanine (pAcF) in C321.ΔA.opt using sfGFP variants with 0, 1, and 3 residues replaced by the UAG codon and confirmed that C321.ΔA.opt maintains nsAA-dependent protein expression (**Fig. 4b**).

**Figure 4.**
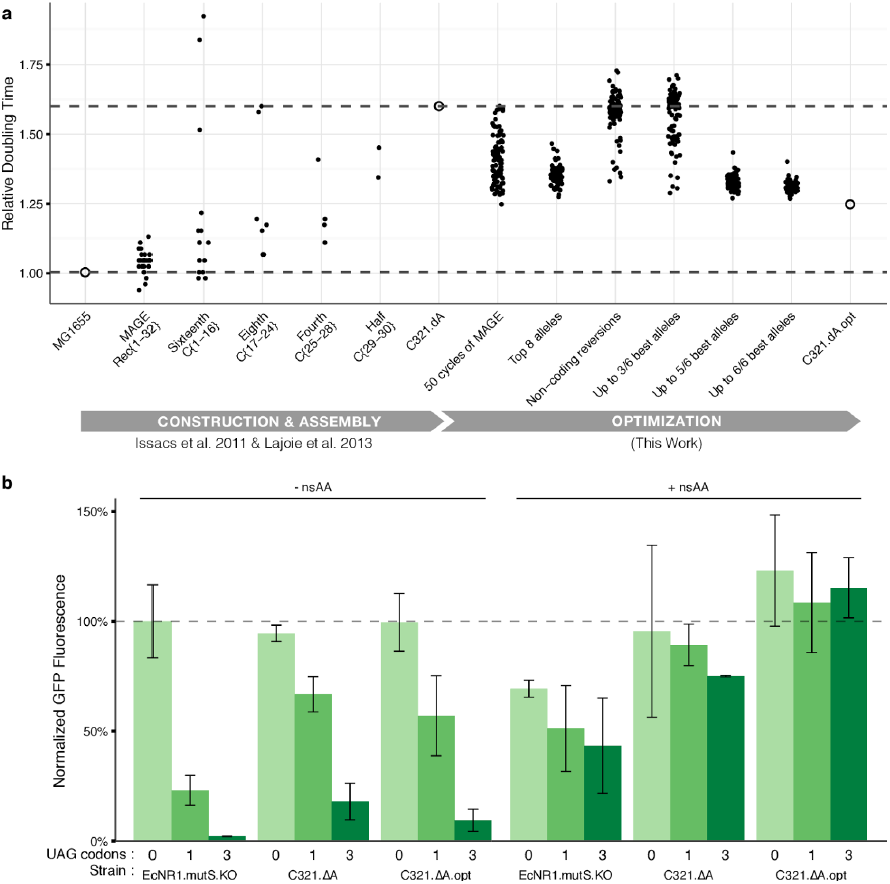
Construction and characterization of final strain C321.ΔA.opt. (**a**) Doubling time of clones isolated during construction and optimization of C321.ΔA. Strain C321, ΔA.opt was constructed in seven cycles of MAGE in batches of up to three cycles separated by MASC-PCR screening to pick clones with the maximum number of alleles converted (see **Online Methods**). The two dotted horizontal lines correspond to the relative doubling times for the original GRO and the wild-type strain. (**b**) Testing nsAA-dependent protein expression using the nsAA p-acetyl-L-phenylalanine (pAcF) in sfGFP variants with 0, 1, or 3 residues replaced with UAG codons. Normalized GFP fluorescence was calculated by taking the ratio of absolute fluorescence to OD600 of cells suspended in Phosphate Buffered Saline (PBS) for each sample and normalizing to the fluorescence ratio of non-recoded strain EcNR1.mutS.KO expressing 0 UAG sfGFP plasmid.

To address the potential benefit of introducing additional changes, and to measure potential interactions among the six alleles identified, we characterized fitness of 359 clones with intermediate genotypes generated during the construction of the final strain (**Fig. 4a**). We applied linear regression with higher order interaction terms (**Fig. 5a**) and observed that combinations of mutations tended to produce diminishing returns ^19^, suggesting that additional beneficial alleles would only contribute marginally to fitness (**Fig. 5b**). We also found evidence of positive epistatic interactions between some alleles (**Fig. 5a**, left), which may not have been identified through singleplex editing strategies. These findings demonstrate the potential use of multiplex genome engineering and predictive modeling for studying epistasis.

**Figure 5.**
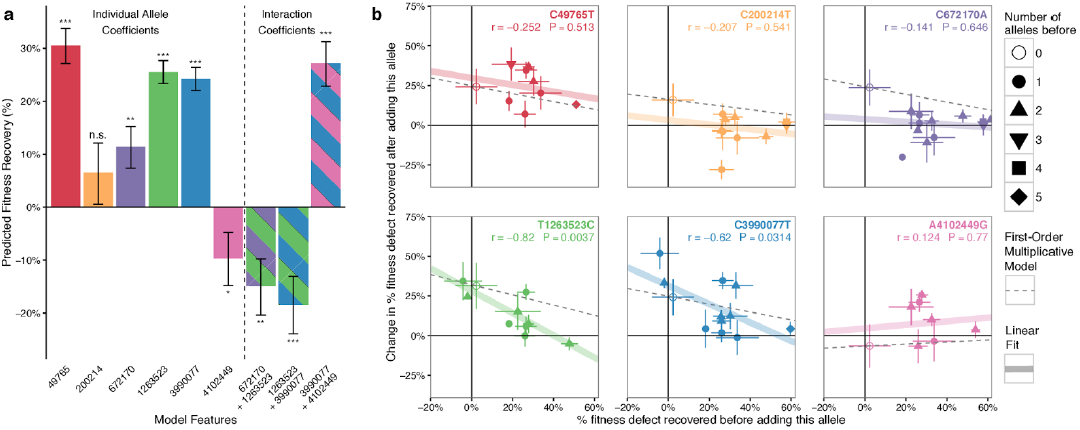
Interactions among top six alleles show evidence of epistasis. Genotypes and fitness measurements were obtained from 359 intermediate clones generated during the the construction of the final strain containing the six best alleles (**Supplementary Table 6**). Each clone was genotyped using MASC-PCR and doubling time was measured during allele validation experiments and final strain construction. (**a**) Individual model coefficients for the top six alleles, as well as three significant interaction terms identified during combinatorial construction. These values are from a linear model with interaction terms between each pair of alleles. The bars signify the standard error of the mean of the model coefficients, and the significance codes for a non-zero effect size are ‘***’: p < 0.001, ‘**’ p < 0.01, p 0. 05, ‘n.s.’ not significant. (**b**) Each data point represents the amount of fitness recovered when adding the allele specified to an identical starting genotype background. Horizontal error bars correspond to the standard deviation of fitness defect among all clones with this starting genotype. Vertical error bars represent the standard deviation of all differences between clones with and without the respective allele. For each plot, the thick colored line represents a simple linear fit through the points, corresponding to the rand p values given in each plot. The dotted line corresponds to the predicted fit for a simple multiplicative model of fitness where the allele always recovers a constant percent of the remaining fitness defect regardless of the background. For all alleles except A4102449G (pink), adding the allele to C321 showed a recovery of the fitness defect (>0 on the y axis), with the percentage of defect recovered decreasing as other alleles are also reverted, consistent with a first-order multiplicative model. In some cases the fitness improvement drops more rapidly than predicted by the multiplicative model (i.e. points below the dotted lines), suggesting diminishing returns epistasis. This is supported by the negative-coefficient interaction terms in panel **a**. In the case of A4102449G there appears to be a negative effect with the mutation alone, but an increase in the presence of other alleles, suggesting possible sign epistasis.

## Discussion

In summary, we used an iterative strategy of multiplex genome engineering and model-guided feature selection to converge on six alleles that together improve the fitness of C321.ΔA by 59%. This method allowed us to quantify the effects of hundreds of individual alleles and then rationally introduce only the minimal set of beneficial genetic changes, reducing unintended effects from additional off-target mutations.

Our approach reveals several problems inherent to using enrichment to rank allelic effect. Our data show that alleles enriched over rounds of selection are not necessarily well-correlated with fitness. Allele enrichment may be affected by differences in recombination efficiency, competition among beneficial alleles through clonal interference, and genetic drift. Iterative targeted editing overcomes these obstacles by allowing the measurement of each allele in many genetic backgrounds, so that linear modeling can quantify its average individual effect.

A similar model-guided approach could be used to augment other multiplex genome modification techniques, including yeast oligo-mediated genome engineering ^20^ or multiplex CRISPR/Cas9-based genome engineering in organisms that support homology-directed double-stranded break repair ^21^. Biosensors tied to selections or screens ^22^ can extend this method to optimize biosynthetic pathways in addition to fitness. Economic improvements in multiplex genome sequencing ^23^ will allow this method to scale to thousands of whole genomes, increasing statistical power and enabling the use of more complex models. Chip-based oligo synthesis enables scaling up the number of genomic sites targeted, allowing thousands of alleles to be tested simultaneously ^24–26^. Finally, making such changes trackable ^27^ for targeted sequencing could further increase the economy, speed, and throughput of this approach.

Efficiently quantifying the effects of many alleles on complex phenotypes is critical not only for tuning synthetic organisms and improving industrially relevant phenotypes, but also for understanding genome architecture. While our method is used here to identify and repair detrimental alleles to improve fitness, it will also enable rapid prototyping of design alternatives and interrogation of genomic design constraints. Iteratively measuring and modeling the effects of large numbers of genomic changes in parallel is a powerful approach to navigate and understand genotype-phenotype landscapes.

## Online Methods

### Media and reagents

All experiments were performed in LB-Lennox (LBL) medium (10 g/L bacto tryptone, 5 g/L sodium chloride, 5 g/L yeast extract) with pH adjusted to 7.45 using 10 M NaOH. LBL agar plates were made from LBL plus 15 g/L Bacto Agar. Selective agents were used at the following concentrations: carbenicillin (50 μg/mL), chloramphenicol (20 μg/mL), gentamycin (5 μg/mL), kanamycin (30 μg/mL), spectinomycin (95 μg/mL), and SDS (0.005%w/v).

### Starting strain

The construction and genotype of engineered *E*. *coli* strain C321.ΔA was previously described in detail ^6^. Here, before improving fitness, we constructed strain *C321.ΔA.mutSfix.KO.tolCfix.Δbla:E* by further modifying C321.ΔA to introduce the following changes: 1) the mutS gene was reinserted into the C321.ΔA strain in its original locus, and MAGE was used to disable the gene by introduction of two internal stop codons and a frameshift, and 2) the carbenicillin-resistance marker bla was swapped for gentamicin resistance marker *aacC1* in the lambda red insertion locus.

### Millstone, software for multiplex genome analysis and engineering

*Millstone* (Goodman et al., submitted) was used throughout the project to rapidly process whole genome sequencing data and identify variants in each sample relative to the reference genome, to explore variant data, and to design oligonucleotides for MAGE. The *Millstone* analysis pipeline takes as input raw FASTQ reads for up to hundreds of clones and a reference genome as Genbank or FASTA format. The software then automates alignment of reads to the reference using the Burrows-Wheeler Aligner (BWA-MEM) followed by single nucleotide variant (SNV) calling using Freebayes. *Millstone* performs variant calling in diploid mode, even for bacterial genomes. This accounts for paralogy in the genome and results in mutation calls being reported as “homozygous alternate” (strong wild-type), “heterozygous” (marginal), or wild-type, along with an “alternate fraction” (AF) field that quantifies the fraction of aligned reads at the locus showing the alternate allele. Marginal calls were inspected on a case-by-case basis using *Millstone’s* JBrowse integration to visualize raw read alignments. *Millstone* provides an interface for exploring and comparing variants across samples. After initial exploration and triage in *Millstone*, we exported the variant report from *Millstone* for further analysis and predictive modeling. In follow-up analysis, we labeled variant calls as ‘marginal’ if the alternate allele fraction was between 0.1 and 0.7.

### Identifying off-target mutations for reversions

For the 50-cycle MAGE experiment, we considered only mutations occurring in regions annotated as coding for a protein or functional RNA. Using Millstone annotations of predicted effect and Keio knock-out collection annotation of essentiality ^17^, we defined three priority categories according to expected effect on fitness (**Supplementary Table 1**). A total of 127 targets were allocated to the three categories to be used for the 50-cycle MAGE experiment.

For a separate experiment, off-target mutations in regulatory regions were selected based on the criteria of predicted regulatory disruption of essential genes and several non-essential genes with particularly strong predicted disruption. Regulatory disruption was determined based on calculating change in 5’ mRNA folding or ribosome binding site (RBS) motif strength for mutations occurring up to 30 bases upstream of a gene. We calculated mRNA folding and ribosome binding site (RBS) motif disruption as described in ^8^. Briefly, the minimum free energy (MFE) of the 5-prime mRNA structure was calculated using Unafol’s `hybrid-ss-min` function ^28^ (T=37 °C), taking the average MFE between windows of RNA (−30, +100) and (−15, +100) relative to the start codon of the gene. Mutations that caused a change in MFE of the mRNA of over 10% relative to the wild-type context were prioritized for testing. To predict RBS disruption, the Salis RBS Calculator ^29^ was provided with sequence starting 20 bases upstream of the gene ATG and including the ATG. Mutations that caused a greater than 10-fold change in predicted expression were included for testing. Finally, we also considered mutations that overlapped promoters of essential genes based on annotations from RegulonDB^30^.

The 20 UAG-reversion targets were chosen when UAGs occurred in essential genes, introduced non-synonymous changes in overlapping genes, or disrupted a predicted regulatory feature as above.

### Multiplex automated genome engineering

Single-stranded DNA oligonucleotides for MAGE were designed using Millstone’s optMAGE integration (https://qithub.com/churchlab/optmaqe). Oligos were designed to be 90 base pairs long with the mutation located at least 20 base pairs away from either end. We used the C321.ΔA reference genome (Genbank accession CP006698.1) for oligo design to avoid inadvertently reverting intentional UAG-to-UAA changes. OptMAGE avoids strong secondary structure (< −12 kcal mol-1) and chooses the sense of the oligo to target the lagging strand of the replication fork ^12^. Phosphorothioate bonds were introduced between the first and second and second and third nucleotides at the 5-prime end of each oligo to inhibit exonuclease degradation ^12^. All DNA oligonucleotides were purchased with standard purification and desalting from Integrated DNA Technologies and dissolved in dH20.

MAGE was performed as described in ^12^, with the following specifications: 1) Cells were grown at 34 °C between cycles. 2) We noted that C321.ΔA exhibits electroporation resistance so a voltage of 2.2 kV (BioRad GenePulser, 2.2 kV, 200 ohms, 25 μF was used for cuvettes with 1mm gap) was chosen based on optimization using a lacZ blue-white screen. 3) Total concentration of the DNA oligonucleotide mixture was 5 μM for all electroporations (i.e., the concentration of each oligo was adjusted depending on how many oligos were included in the pool).

The 50-cycle MAGE experiment was carried out in three lineages, with oligo pool sizes of 26, 49, and 127 consisting of oligos from priority categories {1}, {1,2}, and {1,2,3}, respectively (Supplementary Table 1). Note that we originally began with just two pools--the top 26 and all 127 oligos--, but after 5 MAGE cycles the lineage exposed to all 127 oligos was branched to have a separate lineage with only the 49 category {1, 2} oligos in order to obtain more enrichment of the higher priority targets. In order to prevent any population from acquiring permanent resistance to recombination, we toggled the dual-selectable marker tolC at recombinations 23, 31, and 26 for the three lineages, respectively, as described in ^31^. Briefly, an oligo introducing an internal stop codon in tolC was included in the recombination, and after at least 5 hours of recovery, cells were selected in media containing colicin E1, which is toxic in *tolC^+^ E. coli*. In the subsequent recombination, an oligo restoring tolC function was included in the pool after which cells were selected in the presence of 0.005% SDS (w/v).

Validation MAGE experiments composed of 10 or fewer oligos were carried out for up to 9 MAGE cycles, as we expected adequate diversity based on previous experience with MAGE efficiency.

### Whole genome sequencing

Genomic DNA (gDNA) preparation for whole genome sequencing of 96 clones (only 87 considered in manuscript because sequencing analysis revealed that 9 cultures were polyclonal) was performed as in ^31^. Briefly, gDNA was prepared by shearing using a Covaris E210 AFA Ultrasonication machine. Illumina libraries were prepared for pooled sequencing as previously described ^32^. Barcoded Illumina adapters were used to barcode each strain in a 96-well plate. All 96 genomes were sequenced together on a single lane of a HiSeq 2500 PE150 (Supplementary Table 8). Alternative inexpensive WGS library preparation methods have since become available ^23^.

WGS data was processed to identify clonal genotypes in *Millstone*. Demultiplexed, .fastq reads were aligned to the MG1655 reference genome. SNVs were reported with *Millstone*, as described above. During analysis, marginal calls were visually confirmed by examining alignments using *Millstone’s* JBrowse integration.

### Multiplex allele-specific colony PCR (MASC-PCR)

MASC-PCR was used to assess successful reversions in validation experiments of <= 10 targeted mutations and typically performed for 96 clones in parallel. The protocol was performed as previously described ^6^. Briefly, two separate PCRs, each interrogating up to 10 positions simultaneously, were performed on each clone to detect whether the C321.ΔA or reverted allele was present at each position. For each position, the two reactions shared a common reverse primer but used distinct forward primers differing in at least one nucleotide at the 3’ end to match the SNV being assayed specifically. Positive and negative controls were included when available to aid in discriminating cases of non-specific amplification.

### Measuring fitness

Fitness was determined from kinetic growth (OD600) on a Biotek H-series plate reader. Cells were grown at 34 °C in 150 μL LBL in a flat-bottom 96-well plate at 300 rpm linear shaking. To achieve consistent cell state before reading, clones were picked from agar plates or glycerol, grown overnight to confluence, passaged 1:100 into fresh media, grown again to mid-log (~3 hours), and passaged 1:100 again before starting the read. OD measurements were recorded at 5 minute intervals until confluence. Doubling times were calculated according to t_double_ = c * ln(2)/m, where c = 5 minutes per time point and m is the maximum slope of ln(OD600). The maximum slope was determined using a sliding window linear regression through 8 contiguous time points (40 minutes) points rather than between two predetermined OD600 values because not all of the growth curves were the same shape or reached the same max OD600. The script used for analyzing doubling time is available at https://github.com/churchlab/analyze_plate_reader_growth

### Predictive modeling of allele causality

Choosing alleles for subsequent validation was framed as a feature selection problem. We used predictive modeling to prioritize features. Both targeted reversions introduced by MAGE and de novo mutations were considered.

For most analyses, we used a first-order multiplicative allele effect model, where each allele (reversion or *de novo* mutation) is represented by a single feature and the fitted coefficient corresponding to that feature represents the allele’s effect on doubling time. To find coefficient values, we fit a linear model where genotypes (WGS or MASC-PCR) predict the logarithm of doubling time. Alleles corresponding to features with the most negative coefficients were selected for validation in smaller sets. An additive model was also tested and yielded similar results, as previously noted by others ^19^.

While we anticipated the possibility of epistatic effects among alleles tested, a first-order model of the 50-cycle MAGE experiment already had 239 features (99 reversions + 140 *de novos* observed at least twice) and 87 samples, so we omitted higher-order interaction terms to avoid overfitting due to model complexity. We discuss implications of this independence assumption and other details of our allele effect modeling strategy in Supplementary Note 1.

Elastic net regularization ^18^, which includes both L1 and L2 regularization penalties, was used in model-fitting. L1 regularization enforces sparsity, capturing the assumption that a handful of alleles will explain a majority of the fitness effect. L2 regularization prevents any one of a subset of highly correlated alleles from dominating the effect of those alleles, balancing the tendency of L1 to drop subsets of highly co-occurring alleles.

Accordingly, the elastic net loss function used was

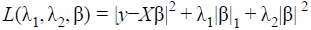

where

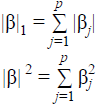

And the coefficients were estimated according to:

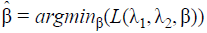

Elastic net regression was performed using the ElasticNetCV module from scikit-learn (Pedregosa et al.). This module introduces the hyperparameters 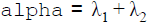 and 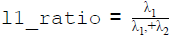 and uses k-fold cross validation (k=5) to identify the best choice of hyperparameters for a given training dataset. We specified the range of `11_ratio` to search over as [.1, .3, .5, .7, .9, .95, .99, 1], which tests with higher resolution near L1-only penalty. This fits our hypothesis that a small number of mutations are responsible for a majority of the fitness effect. For alpha, we followed the default of allowing scikit-learn to search over 100 `alpha` values automatically computed based on `11_ratio`.

To avoid overfitting due to the undersampled nature of the data in the 50-cycle MAGE experiment, we performed 100 repetitions of scikit-learn’s cross-validated elastic net regression procedure, and for each repetition, we randomly held-out 15 samples that could be used to evaluate the model fit by that iteration. The average model coefficient for each allele was then calculated across all 100 repetitions. Only model coefficients with a negative value (some putative fitness improvement) were considered in a second round of 100 repeats of cross-validated elastic net regression, again with 15 samples held-out in each repeat to evaluate the model fit. The average coefficient values over this second set of 100 repetitions were used to determine the top alleles for experimental validation in a 9-cycle MAGE experiment.

To evaluate the results of the 9-cycle MAGE validation experiments, we used unregularized linear regression. With <= 10 parameters and ~90 clones, only a single iteration of cross-validated regression applied to the full dataset was required to assign predicted effects without requiring the testing of individual alleles.

### Final strain construction

C321.ΔA.opt was constructed by adding the six alleles identified by the optimization workflow (**Supplementary Table 6**) to C321.ΔA.mutSfix.KO.tolCfix.Δbla:E. A total of seven cycles of MAGE were required, with a MASC-PCR screening step every three cycles to select a clone with the best genotype so far (**Fig. 3a**), minimizing the total number of cycles required. Three cycles of MAGE were performed using oligos targeting all six alleles. Ninety-six clones were screened by MASC-PCR, and one clone with 3/6 alleles (C49765T, T1263523C, A4102449G) was chosen for the next round of MAGE. Three more rounds of MAGE were performed on top of the clone with 3/6 alleles using only the three remaining oligos. MASC-PCR identified a clone with 5/6 alleles (C49765T, C200214T, C672170A, T1263523C, A4102449G). One more round of MAGE was performed using the remaining oligo and a clone with all six alleles was obtained. Additional off-target mutations acquired during construction as identified by whole genome sequencing of the final clone are listed in **Supplementary Table 7**.

### Characterizing non-standard amino acid incorporation

nsAA incorporation was measured as previously described ^6^. 1-UAG-sfGFP, and 3-UAG-sfGFP reporters were produced by PCR mutagenesis from sfGFP (**Supplementary Note 3**), and isothermal assembly was used to clone 0-UAG-sfGFP (unmodified sfGFP), 1-UAG-sfGFP, and 3-UAG-sfGFP into the pZE21 vector backbone ^33^. We used the pEVOL-pAcF plasmid to incorporate the non-standard amino acid p-acetyl-L-phenylalanine. Reagents were used at the following concentrations: anhydrotetracycline (30 ng/μL), L-arabinose (0.2% w/v), pAcF (1 mM).

## Acknowledgements

We thanks members of the Church Lab for comments. Funding for this work was provided by U.S Department of Energy grant DE-FG02-02ER63445. G.K. and M.J.L. were supported by DOD NDSEG Fellowships. D.B.G. and G.T.F. were supported by NSF Graduate Research Fellowships. Computational resources for this work were provided by the AWS Cloud Credits for Research Program.

## Author contributions

G.K., M.J.L., and D.B.G. designed the study. G.K., M.J.L., G.T.F., and M.M.L. designed and performed experiments. N.R. assisted with whole genome sequencing. G.K., D.B.G., and G.T.F. performed data analysis. G.K., G.T.F., D.B.G., J.A., and M.J.L. wrote the manuscript, and all authors contributed to editing of the manuscript. G.M.C. supervised the project.

## Supplementary Information

**Supplementary Figure 1.**
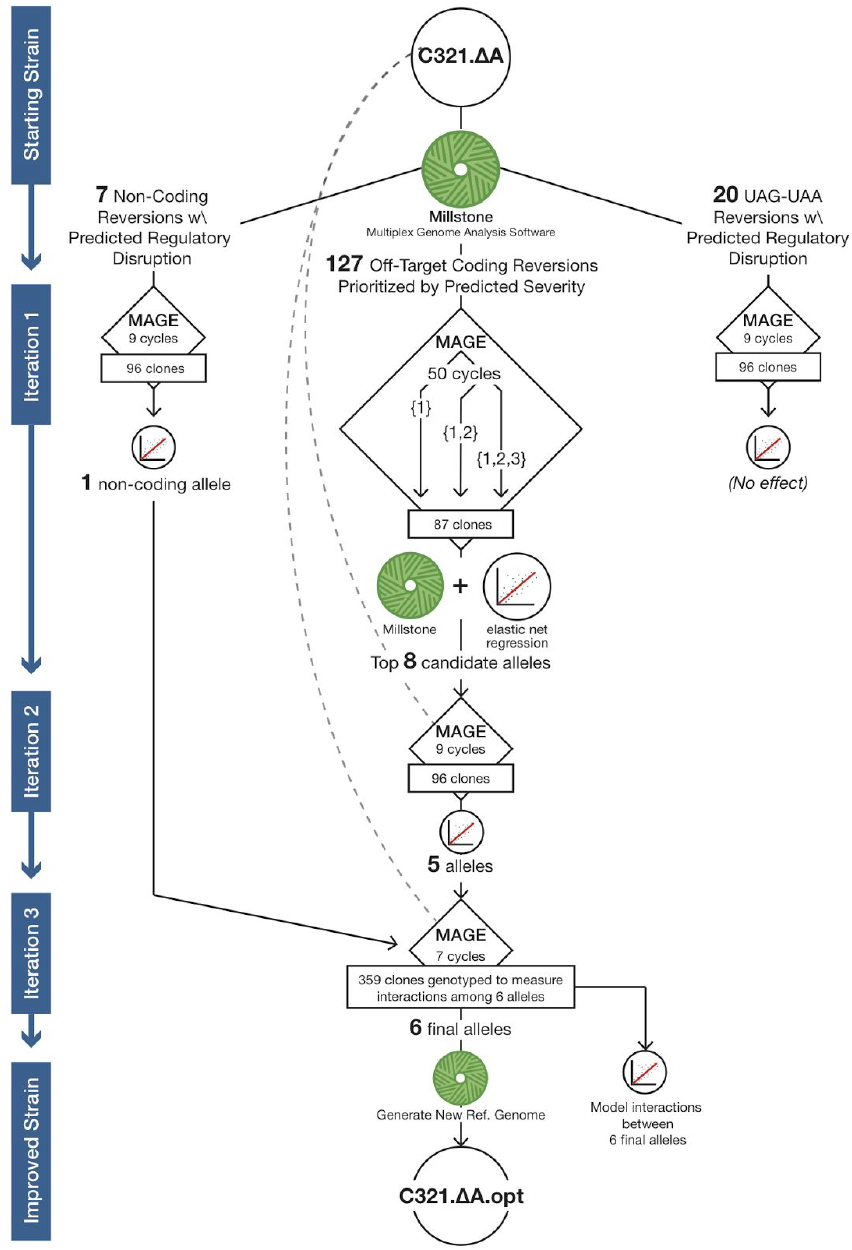
Detailed Experimental Workflow. Depiction of the specific steps used to identify the six alleles that optimized the fitness of C321.ΔA. *Millstone* (Goodman et al., submitted) was used to annotate mutations in C321.ΔA. 127 prioritized coding mutations were tested in C321.ΔA over 50 cycles of MAGE in 3 lineages. Eighty-seven clones were genotyped by whole genome sequencing and annotated using *Millstone*, and their doubling times were measured. Modeling by multiple linear regression identified 8 alleles for subsequent validation. After the second iteration, 5 alleles were chosen, with 3 alleles having a significant linear model coefficient and 2 more reversions having subtle effects. In a parallel experiment, a small pool of 7 non-coding mutations was tested, and modeling identified one allele that was found to have a strong effect. In another parallel experiment, 20 UAA-to-UAG reversions were tested on a C321 background with prfA still present, but no reversions were found to affect fitness. The top 6 alleles were combined in a final optimized strain, and clones with intermediate combinations of alleles were used to characterize their interaction effects (**Supplementary Fig. 5**).

**Supplementary Figure 2.**
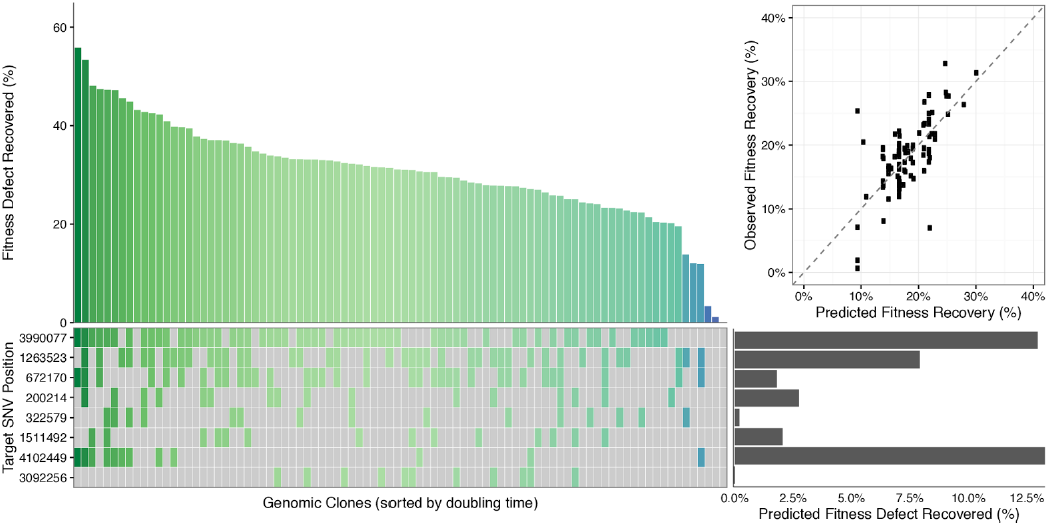
Empirical testing to validate top eight alleles from 50-cycle MAGE experiment. The top eight alleles (**Supplementary Table 3**) were tested in the original C321.ΔA background using nine cycles of MAGE. We selected 96 clones from the final population and measured doubling times and performed MASC-PCR to assess genotypes. Clones are sorted by fitness on the x axis and alleles are listed on the y axis in order of enrichment. Linear modeling revealed a strong predicted effect for reversions T1263523C and A4102449G and *de novo* mutation in C3990077T, with weaker predicted effects for reversions C672170T and C200214T and *de novo* mutation T1511492C. For construction of the final strain, we chose to keep the three high-predicted-effect alleles (T1263523C, A4102449G, C3990077T) and the two weak-predicted-effect reversions (C672170T, C200214T), but we omitted the three weak-effect *de novo* mutations.

**Supplementary Figure 3.**
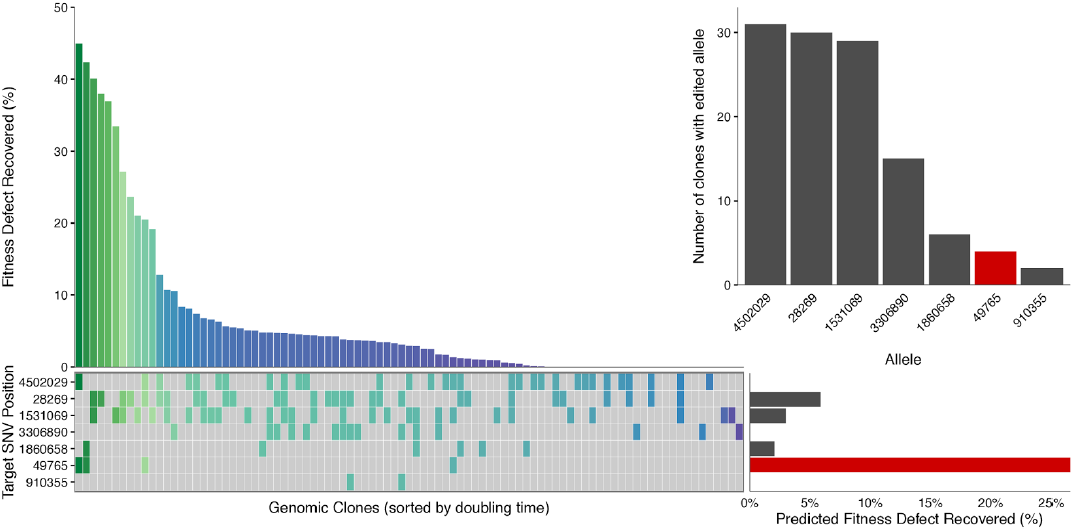
Empirical testing identifies high-effect non-coding mutation. Genotypes and fitness from testing a set of seven non-coding mutations (**Supplementary Table 4**). Upper right: the top model-selected allele is not apparent from enrichment alone, and it is later experimentally validated to have a significant effect (**Figure 4**).

## Supplementary Notes

### Supplementary Note 1

#### Further discussion of allele effect modeling and feature selection

Adaptive laboratory evolution (ALE) typically uses enrichment of mutated genes observed across replicate lineages evolved in parallel to select meaningful features ^34,35^. We selected linear modeling as an alternative to enrichment as initial tests suggested that it was a better method for predicting SNPs that recovered fitness in our experiments. For some alleles, high model coefficients corresponded to high levels of enrichment. For example, the reversion of mutation T1263523C had the highest enrichment after 50 cycles of MAGE (occurring in 78 out of 87 clones) and was also selected by modeling for validation, eventually being verified to confer fitness improvement (**Fig. 2**). On the other hand, when testing the pool of seven non-coding reversions, the single allele selected by the model C49765T (also later experimentally validated) occurred in only 4 out of 96 clones. Meanwhile, three other alleles occurred in over 25 out of 96 clones, but they were not predicted to have a strong effect by linear modeling (**Supplementary Fig. 4**). There were many other cases where linear regression assigned low coefficients to alleles that were highly enriched. The discrepancy between enrichment and model-predicted effect may be due to differences in MAGE oligonucleotide recombination frequency ^6,36^, insufficient time for mutations to achieve enrichment, or stochastic enrichment of passenger mutations in a lineage during MAGE cycling. Altogether, linear modeling provided a more robust strategy of predicting fitness effect for individual alleles.

An important consideration with linear modeling is whether to include higher-order interaction terms. For our 50-cycle MAGE experiment, we made an assumption that independent allele effects would dominate relative to complex epistatic effects and included only first-order terms in the model. For validation in pools of <= 10 oligos, we typically selected 96 clones and experimented with second- and higher-order models, but also found that the first-order model was typically sufficiently informative of allele effect.

To investigate how higher-order model terms can inform interpretation of epistatic effects, we assembled a dataset of 359 intermediate genotyped clones obtained from validation experiments or from screening during construction of the final strain with all six top alleles (**Supplementary Fig. 5**). Interestingly, linear modeling with second-order interaction terms indicated evidence of possible diminishing returns epistasis among certain alleles ^37^ and also a possible positive epistasis effect between A4102449G and C3990077T. Alleles that contribute to fitness through a positive epistasis effect could be lost during validation of small numbers of alleles, supporting our validation of high impact alleles in pools.

Even with targeted engineering by MAGE, de novo mutations can play a role in fitness improvement. The mismatch repair-deficient context in which this study was conducted elevates the background mutation rate >100-fold 38 and resulted in the accumulation of four de novo mutations for every reversion (**Supplemental Fig. 2f**). We considered de *novo* mutations in modeling experiment data, but omitted any de *novo* mutation that was never observed in more than one clone, reducing the number of features corresponding to de *novo* mutations from 1329 to 135. Linear regression of data obtained over 50-cycles of MAGE identified four *de novo* mutations with a putative effect. Validation of these alleles determined that three of these were false positives or only beneficial in a specific context. The fourth *de novo* C3990077T, however, showed a strong effect upon validation, demonstrating that linear regression can be an effective strategy for identifying causal *de novo* mutations and may be generally applicable in laboratory evolution studies.

Additional features beyond allele occurrence can be added to the linear model such as terms that capture prior expectation of an allele’s effect. For application to ALE, mutations could be merged according to affected gene and its interacting partners. Higher order terms can be iteratively introduce as the candidate feature set is pruned.

### Supplementary Note 2

#### Discussion of alleles chosen to construct final strain

The final strain was constructed by introducing six mutations into the starting C321.ΔA background: five alleles reverted to their MG1655 starting point and one de novo mutation not previously present in MG1655 (**Supplementary Table 6**).

Four of five reversions were coding mutations in essential genes prioritized in the highest category in the 50-cycle MAGE experiment, supporting the strength of the initial prioritization method. The fifth reversion was identified in screening noncoding off-target mutations predicted to disrupted gene regulation, demonstrating computational prediction of regulatory disruption as an important strategy in tuning organisms ^8^. The sixth mutation

We characterized intermediate genotypes created while constructing the final strain (**Supplementary Fig. 5**) and determined that three of the mutations (reversions of C49765T, T1263523C and de novo C3990077T) had especially strong individual effects. Two reversions (C672170T, C200214T) had weaker individual effects that diminished in backgrounds with multiple mutations (**Supplementary Fig. 5b**). The last reversion A4102449G did not have a strong effect alone, and may even have been slightly detrimental alone, but appeared to provide a benefit in the presence of C3990077T (**Supplementary Fig. 5a**).

We suspected that the *de novo cyaA* mutation (C3990077T) is a beneficial suppressor in the C321.ΔA background but not the non-recoded background. Testing the mutation in EcNR1.mutS.KO revealed a minor detrimental effect on fitness, increasing doubling time by 2.94% (p=0.002; one-tailed t-test).

### Supplementary Note 3

#### Nucleotide sequences for 0-UAG-sfGFP, 1-UAG-sfGFP, 3-UAG-sfGFP used for characterizing nsAA incorporation

~~~
>0-UAG-sfGFP
ATGCATCACCACCATCATCACAAAGGTGAAGAACTGTTTACCGGCGTTGTTCCGATCCTGGTTGAACTGGACGGTGACGTGAACGGTCATAAATTCTCCGTACGTGGTGAAGGTGAGGGTGACGCGACCAACGGTAAGCTGACTCTGAAATTCATCTGCACCACCGGCAAACTGCCGGTTCCGTGGCCGACGCTGGTTACGACCCTGACCTACGGTGTTCAGTGCTTCGCGCGTTACCCGGACCATATGAAGCAGCACGACTTCTTCAAATCTGCGATGCCGGAAGGTTACGTTCAGGAACGTACCATCTCTTTCAAAGACGACGGTACCTACAAAACCCGTGCGGAAGTTAAATTCGAAGGCGACACCCTGGTTAATCGTATCGAACTGAAAGGTATCGACTTCAAGGA AG ACG GC AATATTCTGG GT C ACAAACT G G AATACAACTT C AACT CT CAC AAT GTTTACAT C ACCG CG G A CAAACAGAAAAATGGTATCAAAGCAAATTTCAAAATCCGTCATAACGTTGAGGACGGCTCTGTACAACTGGCGGACCACTACCAACAAAACACCCCGATTGGTGACGGTCCGGTCCTGCTGCCGGACAACCATTACCTGTCTACCCAGTCTGTTCTGTCTAAAGACCCGAACGAAAAACGTGACCACATGGTTCTGCTGGAATTCGTTACCGCAGCGGGTATCACCCACGGTATGGACGAGCTGTATTAA
~~~

~~~
>1-UAG-sfGFP
ATGCATCACCACCATCATCACAAAGGTGAAGAACTGTTTACCGGCGTTGTTCCGATCCTGGTTGAACTGGACGGTGACGTGAACGGTCATAAATTCTCCGTACGTGGTGAAGGTGAGGGTGACGCGACCAACGGTAAGCTGACTCTGAAATTCATCTGCACCACCGGCAAACTGCCGGTTCCGTGGCCGACGCTGGTTACGACCCTGACCTACGGTGTTCAGTGCTTCGCGCGTTACCCGGACCATATGAAGCAGCACGACTTCTTCAAATCTGCGATGCCGGAAGGTTACGTTCAGGAACGTACCATCTCTTTCAAAGACGACGGTACCTACAAAACCCGTGCGGAAGTTAAATTCGAAGGCGACACCCTGGTTAATCGTATCGAACTGAAAGGTATCGACTTCAAGGAAGACGGCAATATTCTGGGTCACAAACTGGAATACAACTTCAACTCTCACAATGTTTAGATCACCGCGGACAAACAGAAAAATGGTATCAAAGCAAATTTCAAAATCCGTCATAACGTTGAGGACGGCTCTGTACAACTGGCGGACCACTACCAACAAAACACCCCGATTGGTGACGGTCCGGTCCTGCTGCCGGACAACCATTACCTGTCTACCCAGTCTGTTCTGTCTAAAGACCCGAACGAAAAACGTGACCACATGGTTCTGCTGGAATTCGTTACCGCAGCGGGTATCACCCACGGTATGGACGAGCTGTATTAA
~~~

~~~
>3-UAG-sfGFP
ATGCATCACCACCATCATCACAAAGGTGAAGAACTGTTTACCGGCGTTGTTCCGATCCTGGTTGAACTGGACGGTGACGTGAACGGTCATAAATTCTCCGTACGTGGTGAAGGTGAGGGTGACGCGACCTAGGGTAAGCTGACTCTGAAATTCATCTGCACCACCGGCAAACTGCCGGTTCCGTGGCCGACGCTGGTTACGACCCTGACCTACGGTGTTCAGTGCTTCGCGCGTTACCCGGACCATATGAAGCAGCACGACTTCTTCAAATCTGCGATGCCGGAAGGTTACGTTCAGGAACGTACCATCTCTTTCAAAGACGACGGTACCTACAAAACCCGTGCGGAAGTTAAATTCGAAGGCGACACCCTGGTTAATCGTATCGAACTGAAAGGTATCGACTTCAAGGAAGACGGCAATATTCTGGGTCACAAACTGGAATACAACTTCAACTCTCACAATGTTTAGATCACCGCGGACAAACAGAAAAATGGTATCAAAGCAAATTTCAAAATCCGTCATAACGTTGAGGACGGCTCTGTACAACTGGCGGACCACTAGCAACAAAACACCCCGATTGGTGACGGTCCGGTCCTGCTGCCGGACAACCATTACCTGTCTACCCAGTCTGTTCTGTCTAAAGACCCGAACGAAAAACGTGACCACATGGTTCTGCTGGAATTCGTTACCGCAGCGGGTATCACCCACGGTATGGACGAGCTGTATTAA
~~~

**Table S1: Prioritized coding reversion categories**

**Table S2: Table of 127 targeted mutations**

**Table S3: Top mutations from MAGE cycling**

**Table S4: List of non-coding mutations tested**

**Table S5: List of amber reversions tested**

**Table S6: List of top six mutations used to create C321.DA.opt**

**Table S7: List of additional mutations in C321.DA.opt**

**Table S8: Cost analysis**

**Table S9: Mutations from whole genome sequencing data for 90 clones from 50-cycle MAGE experiment**

